# Adult expression of the cell adhesion protein Fasciclin 3 is required for the maintenance of adult olfactory interneurons

**DOI:** 10.1101/2024.03.24.586496

**Authors:** Aarya Vaikakkara Chithran, Douglas W. Allan, Timothy P. O’Connor

## Abstract

The proper functioning of the nervous system is dependent on the establishment and maintenance of intricate networks of neurons that form functional neural circuits. Once neural circuits are assembled during development, a distinct set of molecular programs is likely required to maintain their connectivity throughout the lifetime of the organism. Here, we demonstrate that Fasciclin 3 (Fas3), an axon guidance cell adhesion protein, is necessary for the maintenance of the olfactory circuit in adult *Drosophila*. We utilized the TARGET system to spatiotemporally knockdown *Fas3* in selected populations of adult neurons. Our findings show that *Fas3* knockdown results in the death of olfactory circuit neurons and reduced survival of adults. We also demonstrated that *Fas3* knockdown activates caspase-3 mediated cell death in olfactory local interneurons, which can be rescued by overexpressing p35, an anti-apoptotic protein. This work adds to the growing set of evidence indicating a critical role for axon guidance proteins in the maintenance of neuronal circuits in adults.

**SUMMARY STATEMENT:** Little is known about the maintenance of adult neural circuits. We show that the continuous expression of Fasciclin 3, a cell adhesion protein involved in axon guidance, is required for neuronal survival in the adult olfactory circuit.

## INTRODUCTION

The proper functioning of the nervous system is determined by the establishment and maintenance of intricate connections between neurons. During nervous system development, billions of neuronal axons navigate to their targets in response to axon guidance cues in the extracellular environment (Tessier-Lavigne and Goodman, 1996). This process involves neuronal outgrowth, navigation and pathfinding that relies on a highly dynamic, fan-shaped, actin-rich structure located at the tip of axons and dendrites called a growth cone (Cajal, 1890). For over 100 years, neuroscientists have researched the nature of the guidance cues underlying these processes and developed an understanding of how these cues signal spatial information to growth cones to direct neural circuit formation (Kolodkin and Tessier-Lavigne, 2011). Many families of guidance cues have been characterized including Semaphorins, Netrins, Slits, Ephrins, morphogens, cell adhesion molecules (CAMs) and cytoskeletal-associated proteins (Yaron and Zheng, 2007).

Numerous studies have supported the idea that CAMs can guide axons by promoting adhesion, stimulating neuronal outgrowth, and functioning as signaling molecules in heterophilic or homophilic combinations (Moreland and Poulain, 2022; Pollerberg et al., 2013; Rutishauser, 1993). The immunoglobulin-like cell adhesion molecules (Ig-CAMs) that belong to the immunoglobulin superfamily and cadherins are the two major classes of CAMs. A role for CAMs in regulating axonal fasciculation and motor neuron guidance was first documented for the Ig-CAM Fasciclin 2 (Fas2) (Harrelson and Goodman, 1988; Lin and Goodman, 1994; Lin et al., 1994). Fas2 is homologous to the neural cell adhesion molecule (N-CAM) in vertebrates and is required for selective axon fasciculation in *Drosophila*. Ig-CAMs are also known to interact with signal transduction pathways during axonal guidance. For example, *Fasciclin 1* (*Fas1*) genetically interacts with *Abelson tyrosine kinase* (*Abl*) with *Fas1/Abl* double mutants exhibiting commissural axon pathfinding defects (Elkins et al., 1990). Fasciclin 3 (Fas3), another Ig-CAM in *Drosophila,* is expressed by RP3 motor neurons and their synaptic targets (muscles 6 and 7) during embryonic development (Patel et al., 1987; Snow et al., 1989). Target recognition between the motor neuron growth cones and muscles is mediated through *Fas3*-dependent homophilic interactions. In addition, RP3 motor neurons form synapses on muscles that ectopically express *Fas3*, confirming that *Fas3* acts as a synaptic recognition cue for these growth cones (Chiba et al., 1995). Additionally, when the *Fas3*-negative aCC and SNa motor neuron growth cones ectopically express *Fas3*, they innervate the *Fas3*-expressing muscles as alternative targets (Kose et al.,1997). However, motor neurons can correctly innervate the target muscles 6 and 7 in *Fas3* null mutants, suggesting redundancy in targeting mechanisms. These studies provide evidence for *Fas3*-dependent homophilic synaptic target recognition in the precise formation of neural circuits.

Once neural circuits are assembled during development, it is presumed that a distinct set of molecular programs is likely required to maintain their connectivity throughout the lifetime of the organism. Nonetheless, many mature neurons continue to express axon guidance proteins long after they have reached their targets and made functional connections. Indeed, the modENCODE high-throughput RNA-seq data confirms that axon guidance cues, including Semaphorins, Plexins, and CAMs (including *Fas3*) continue to be expressed in the adult *Drosophila* nervous system (Graveley et al., 2011). We have recently shown that members of the Semaphorin family of guidance cues have critical functions in the mature nervous system and are essential for maintaining neuronal survival, adult motility and longevity (Vaikakkara Chithran et al., 2023). Although this previous work showed that the expression of *Fas3* in adults is also critical for survival, the role of *Fas3* in the maintenance of adult neural circuits has not been examined. In this study, we utilize the well-characterized *Drosophila* olfactory circuit to explore the importance of stable expression of *Fas3* for neural circuit maintenance in adults.

In *Drosophila*, the olfactory lobe (OL) is composed of 56 neuropil regions, known as glomeruli (Laissue et al., 1999, Tanaka et al., 2012), each of which receives input from a specific population of olfactory sensory neurons (OSNs) that typically express a single type of odorant receptor (Couto et al., 2005; Fishilevich and Vosshall, 2005). Within each glomerulus, OSNs establish synapses with projection neurons (PNs), which serve as output neurons of the OL and can be compared to the mitral/tufted cells in mammals (Imai et al., 2010; Lledo et al., 2005). OSNs also make connections with local interneurons (LNs) that have arborizations confined within the OL, similar to interneurons found in the olfactory bulb of mammals, such as granule and periglomerular cells (Lledo et al., 2005; Stocker et al., 1990; Tanaka et al., 2012). Furthermore, PNs establish feedback connections with LNs (Liu and Wilson, 2013; Sudhakaran et al., 2012; Tanaka et al., 2009). LNs play multiple roles in shaping the output of the OL. One important function of LNs is their involvement in olfactory habituation, a phenomenon characterized by a reduced behavioral response to repeated or continuous exposure to an odorant (Twick et al., 2014). This reduced response is attributed to increased inhibitory input from LNs to odor-selective PNs (Das et al., 2011; Larkin et al., 2010; Sadanandappa et al., 2013; Sudhakaran et al., 2012). Interestingly, a genetic screen identified mutations in several CAMs, including *Fas3*, which disrupted olfactory habituation (Eddison et al., 2012), suggesting an important role in the proper development of the olfactory circuit. However, the role of *Fas3* in the normal functioning of olfactory neurons in adults has not been examined. In this study, we present evidence demonstrating the essential role of continuous *Fas3* expression in maintaining the olfactory circuitry in adult flies. Specifically, we reveal that *Fas3* promotes cell survival in LNs, suggesting a critical function for CAMs in the maintenance of adult neurons.

## RESULTS

### *Fas3* is widely expressed in the adult *Drosophila* central nervous system

Using a *Fas3::GFP* protein trap allele combined with Bruchpilot (Brp) immunostaining we examined the expression and localization of *Fas3* in the adult *Drosophila* central nervous system (CNS). *Fas3* is prominently localized in the olfactory lobes (yellow box in Fig. 1A), with lower levels observed in the optic lobes (white box in Fig. 1A), suboesophageal ganglion (yellow arrow in Fig. 1A), and a subset of neuronal populations in the ventral nerve cord (white arrows in Fig. 1A). This expression and localization were also confirmed by Fas3 immunoreactivity (Fig. 1B) and *Fas3-Gal4* driven expression of nuclear GFP (Fig. 1C). Within the olfactory circuit, it is likely that Fas3 is expressed in a subset of neurons whose cell bodies lie adjacent the olfactory lobe (as observed by the *UAS-nuclear GFP* expression driven by *Fas3-Gal4* in Fig. 1C) and project their dendrites into the olfactory lobe (as seen by the strong expression of *Fas3-GFP* and anti-Fas3 labelling in the glomeruli of the olfactory lobe in Fig. 1A, B).

**Fig. 1.**
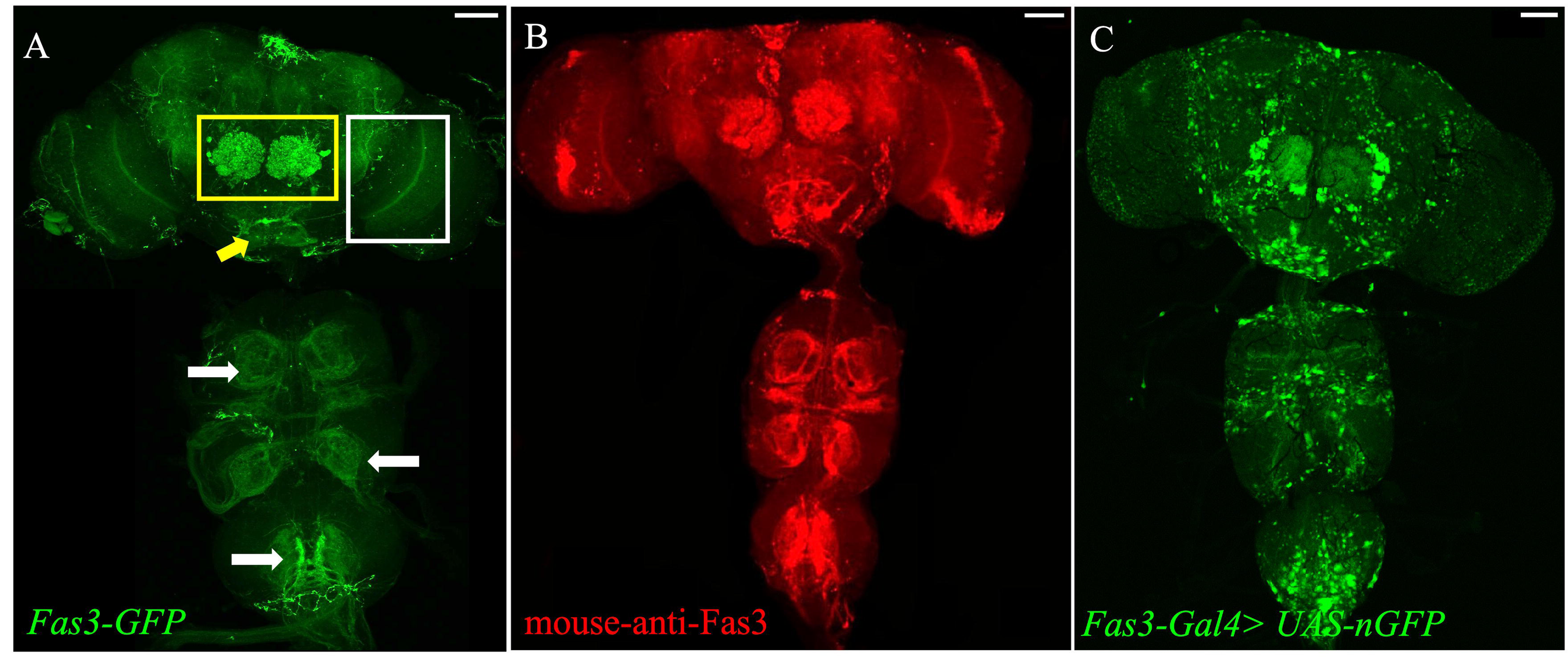
*Fas3* is widely expressed in the adult *Drosophila* CNS. *Fas3* is expressed in the adult *Drosophila* olfactory lobes (yellow box), optic lobes (white box), suboesophageal ganglion (yellow arrow) and ventral nerve cord (white arrows) as shown by (A) *Fas3-GFP* fusion protein expression, (B) immunostaining using a mouse-anti-Fas3 antibody and (C) *UAS-nuclear GFP* (*nGFP*) expression driven by *Fas3-Gal4*. The scale bars represent 50 μm. Images shown are maximum intensity projections. Genotypes: (A) *w*; Fas3-GFP* (B) *Oregon R* (C) *w; Fas3-Gal4/+; UAS-nGFP/+.* See Supplementary Tables S1 and S4 for the additional details on the fly lines.

### *Fas3-Gal4* mediated *Fas3* knockdown reduces adult *Drosophila* survival

To examine if *Fas3* expression in adults is essential for survival, we used the TARGET system (*Fas3-Gal4* with *tub-Gal80^ts^*) to knockdown the expression of *Fas3* only in adult *Drosophila* in all *Fas3*-expressing cells (Fig. 2A) and monitored the survival of adult flies daily until day 22 post eclosion. We picked male flies for all the experiments. In the survival assay, we employed two controls; an isogenic host strain as a genetic background control for the GD RNAi library and a *UAS-dsRNA-GFP* control, both of which yielded similar results. On day 17, there was a significant reduction in survival in the *Fas3* knockdown flies, with only 87% of the knockdown flies surviving compared to 97% of the control flies. This impact on survival increased with age; by day 22, the survival rate of *Fas3* knockdown flies dropped to 47%, while 97% of the control flies survived (Fig. 2B), confirming that continued expression of *Fas3* in the adult is essential for longevity.

**Fig. 2.**
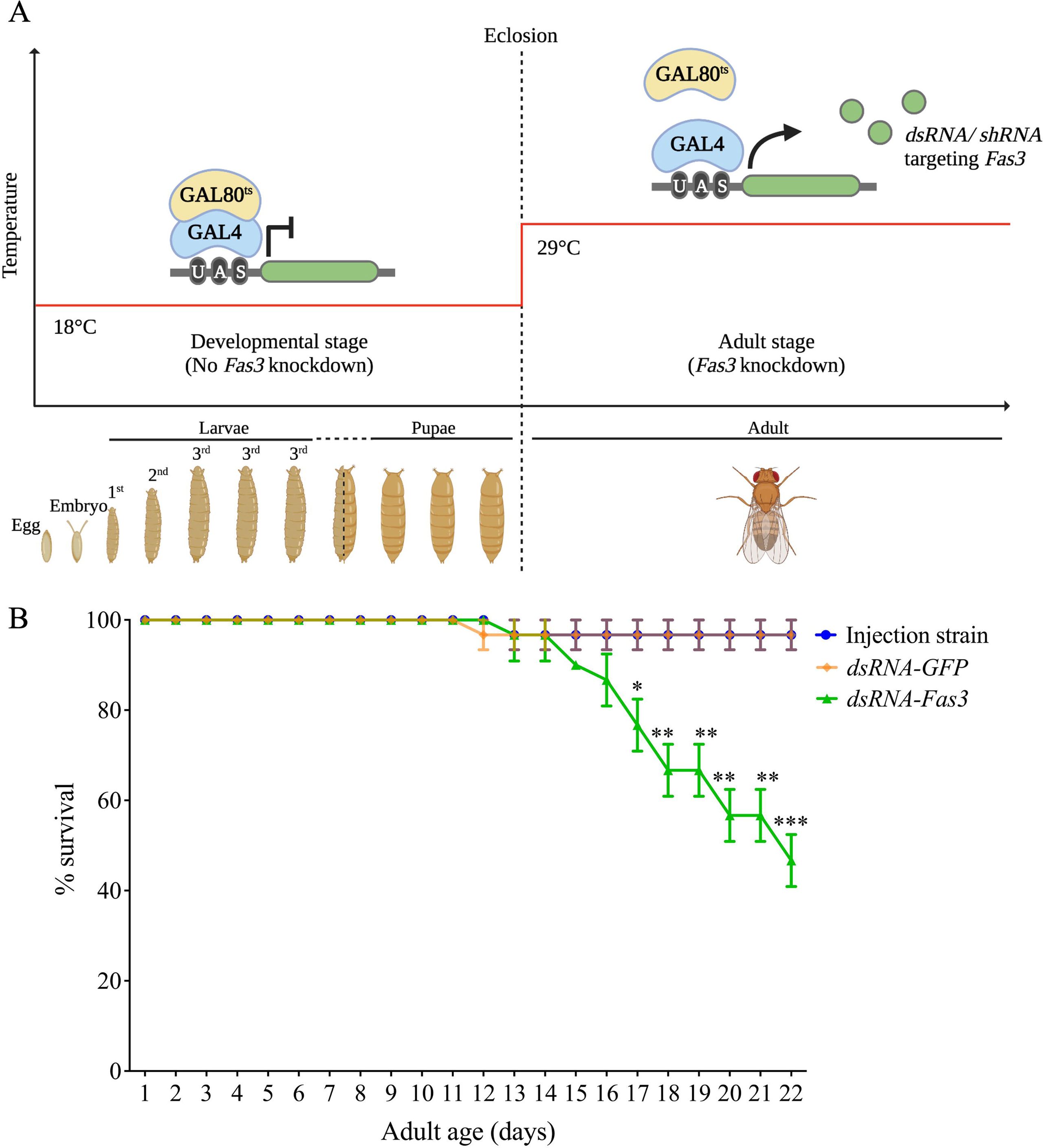
*Fas3-Gal4* mediated *Fas3* knockdown reduces adult *Drosophila* survival from day 17. (A) A schematic of the TARGET system utilized to restrict the *Fas3* knockdown spatiotemporally to adult neurons. After eclosion, flies are moved to the permissive temperature (29°C) to initiate *dsRNA/ shRNA-Fas3* expression. (B) Knockdown of *Fas3* specifically in *Fas3*-expressing cells reduces adult survival. ‘*’ indicates the time points when the survival of *Fas3* knockdowns is significantly different from that of both the injection strain control and the GFP control. Data shown are mean ± SEM. ‘*’ denotes P ≤ 0.05, ‘**’ denotes P ≤ 0.01, and ‘***’ denotes P ≤ 0.001 (two-way ANOVA and Tukey’s multiple comparison tests; n=30). Genotypes: ‘Injection strain’ is *UAS-Dicer2; Fas3-Gal4/+; tubGal80^ts^/+. ‘*dsRNA-GFP’ is *UAS-Dicer2; Fas3-Gal4/ UAS-dsRNA-GFP; tubGal80^ts^/+. ‘*dsRNA-Fas3’ is *UAS-Dicer2; Fas3-Gal4/ UAS-dsRNA-Fas3 (#1); tubGal80^ts^/+.* See Supplementary Tables S1-S3 for the additional details on the fly lines.

### *Fas3* is expressed in a subset of olfactory local interneurons

Due to high *Fas3* levels in the adult olfactory lobes (as shown in the yellow box in Fig. 1A) and the well-characterized circuitry of the adult *Drosophila* olfactory system (Fig. 3A), we examined the impact of *Fas3* knockdown on the maintenance of this circuit. To visualize the precise location of *Fas3* expressing neurons and their processes, we used *Fas3-Gal4* to drive *UASCD8:: GFP*. We observed high expression in neurons around the adult olfactory lobe whose cell bodies lie adjacent to the OL (Fig. 3C). Among the three neuronal populations of the olfactory circuit (OSNs, PNs, LNs), the OSNs have their cell bodies located in the periphery and the PNs send their axonal projections into higher brain regions. However, we observed *Fas3* positive cell bodies right outside the olfactory lobe (Fig 1C) and after extensive examination we could not identify any projections directed to any higher brain regions (Fig 1A, B) suggesting that the *Fas3*-expressing neurons only send local projections and are likely to be LNs.

**Fig. 3.**
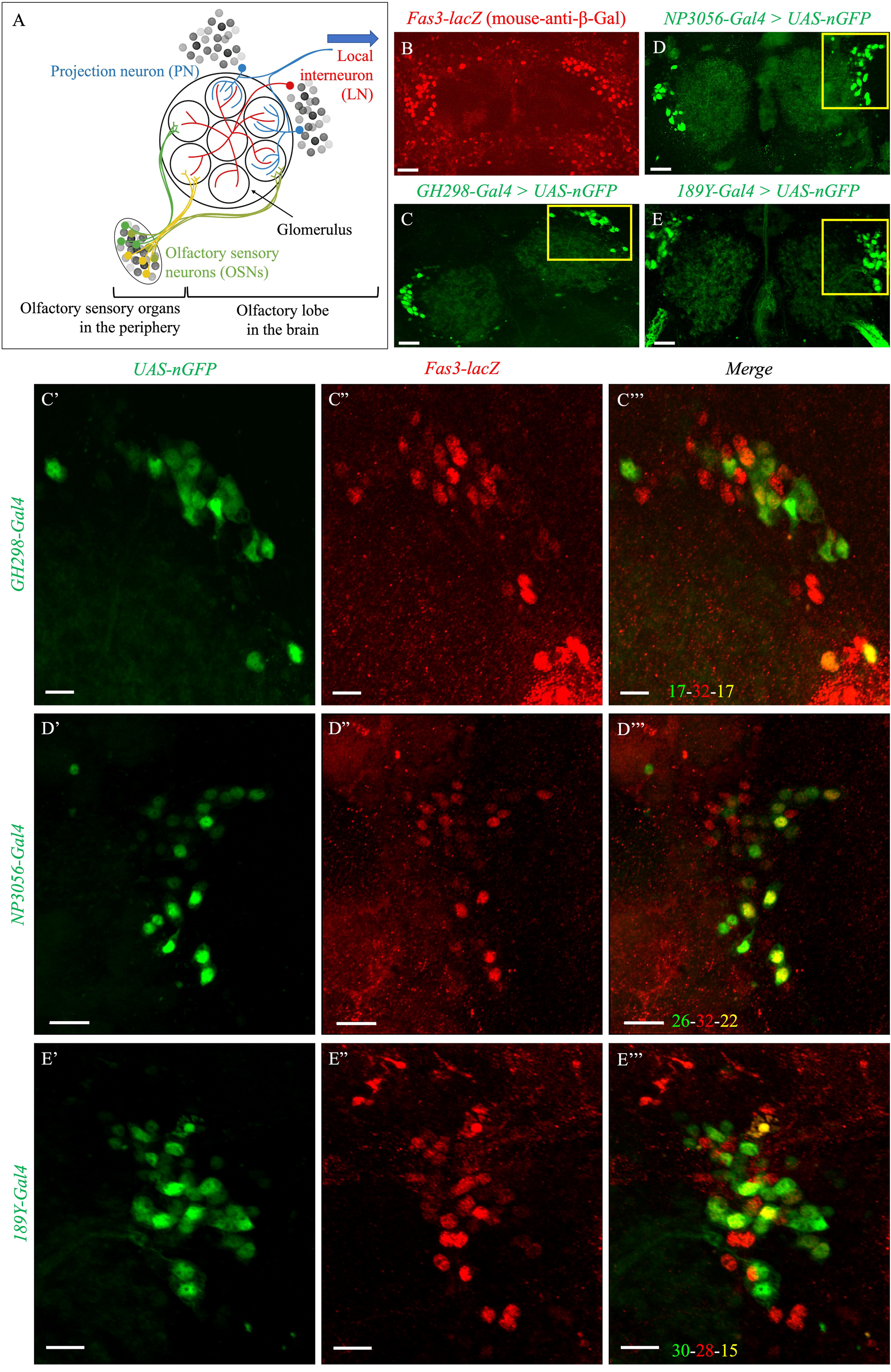
*Fas3* is expressed in olfactory local interneurons. (A) A schematic of the adult *Drosophila* olfactory circuit, adapted from Seki et al., 2010. The axons of OSNs from the periphery project into distinct neuropil regions called glomeruli in the olfactory lobe. Within a given glomerulus, OSNs (yellow and green neurons) form synapses with PNs (blue) and LNs (red). The cell bodies of PNs and LNs lie outside the olfactory lobes. PNs extend their dendrites within specific glomeruli and send their axonal projections to higher centers (blue arrow) in the brain, whereas LNs send their projections locally, within the olfactory lobe. (B) Nuclear lacZ expression in *Fas3*-positive cells shows that the cell bodies are grouped around the adult olfactory lobes. (C-E) The Gal4 drivers *GH298-Gal4*, *NP3056-Gal4* and *189Y-Gal4* are expressed in a subset of LNs. (C’-E’’’) *Fas3* is co-expressed in a subset of the *LN-Gal4* neurons. The numbers shown in the bottom left corner of each merged panel indicate the number of cells labelled by *LN-Gal4> UAS-nGFP* (green), *Fas3-lacZ* (red) and both (yellow). (C’-C’’’) All the neurons labelled by *GH298-Gal4* were also labelled by *Fas3-lacZ* (n=10). (D’-D’’’) Approximately 85% of the neurons labelled by *NP3056-Gal4* also showed *Fas3-lacZ* expression (n=10). (E’-E’’’) Roughly 50% of the *189Y-Gal4* neurons co-expressed *Fas3-lacZ* (n=10). The scale bars represent 20 μm in B-E and 10 μm in C’-E’’’. Images shown are maximum intensity projections. Genotypes: (B) *Fas3-lacZ, CyO/ In(2LR)Gla, wg^Gla-1^.* (C-C’’’) *w; Fas3-lacZ, CyO/+; UAS-nGFP/ GH298-Gal4.* (D-D’’’) *w; Fas3-lacZ, CyO/+; UAS-nGFP/ NP3056-Gal4.* (E-E’’’) *w; Fas3-lacZ, CyO/ 189Y-Gal4; UAS-nGFP/+.* See Supplementary Tables S1 and S4 for the additional details on the fly lines.

To confirm Fas3 expression in LNs, we examined co-labelling of *Fas3-nlacZ* reporter (Fig. 3D) with UAS-nGFP driven by one of three *LN-Gal4* lines, *GH298-Gal4* (Fig. 3E), *NP3056-Gal4* (Fig. 3F) and *189Y-Gal4* (Fig. 3G) (Chou et al., 2010; Liou et al., 2018). Almost all *GH298-Gal4* neurons were co-labelled by *Fas3-lacZ* (Fig. 3E’). Also, 85% of *NP3056-Gal4* neurons (Fig. 3F’) and 50% of *189Y-Gal4* neurons (Fig. 3G’) were co-labelled with *Fas3-lacZ*. This confirmed that *Fas3* is expressed in subsets of olfactory LNs.

### *Fas3* knockdown in the adult results in the death of olfactory circuit neurons

In order to examine the importance of *Fas3* expression in neural circuit maintenance, understand why Fas3 may be essential for adult survival, and due to its strong expression in the olfactory lobes, we decided to examine the impact of *Fas3* knockdown specifically on this circuit. As we observed a significant reduction in the survival of adult flies from day 17 post eclosion (Fig. 2B), we focused our analysis on circuit structure asked whether there was an impact on neuronal integrity before this time point. Specifically, we examined LNs and their projections at day 14, two weeks following initiation of *Fas3* knockdown. Using *Fas3-Gal4*, we drove *UAS-dsRNA-Fas3* with *UAS-CD8::GFP* or as a control we drove *UAS-CD8::GFP* alone. In the controls, we observed the normal distribution of CD8::GFP neuronal cell bodies located immediately adjacent to the olfactory lobes (indicated by white arrows in Fig. 4A). In contrast, in Fas3 knockdowns, we observed a striking, significant loss of these neurons (indicated by white arrows in Fig. 4B). This neuronal loss phenotype was observed with multiple *dsRNA-Fas3* lines (Fig. S1). Fas3 antibody staining confirmed that *Fas3* was severely reduced by day 14 (compare Fig. 4A’, B’). Next, we observed an increase in anti-Caspase 3 immunolabelling as early as day 7, especially in the region surrounding the olfactory lobes (indicated by white arrows in Fig. 4A”, B”). This suggested that loss of olfactory circuit neurons observed in *Fas3* knockdowns is due to apoptotic cell death. It is interesting to note that the region of increased caspase3 appears to extend beyond Fas3 knockdown, suggesting possible apoptosis in other cells immediately surrounding the Fas3 expressing neurons. Further experiments will be required to examine the possible mechanisms underlying this cell death.

**Fig. 4.**
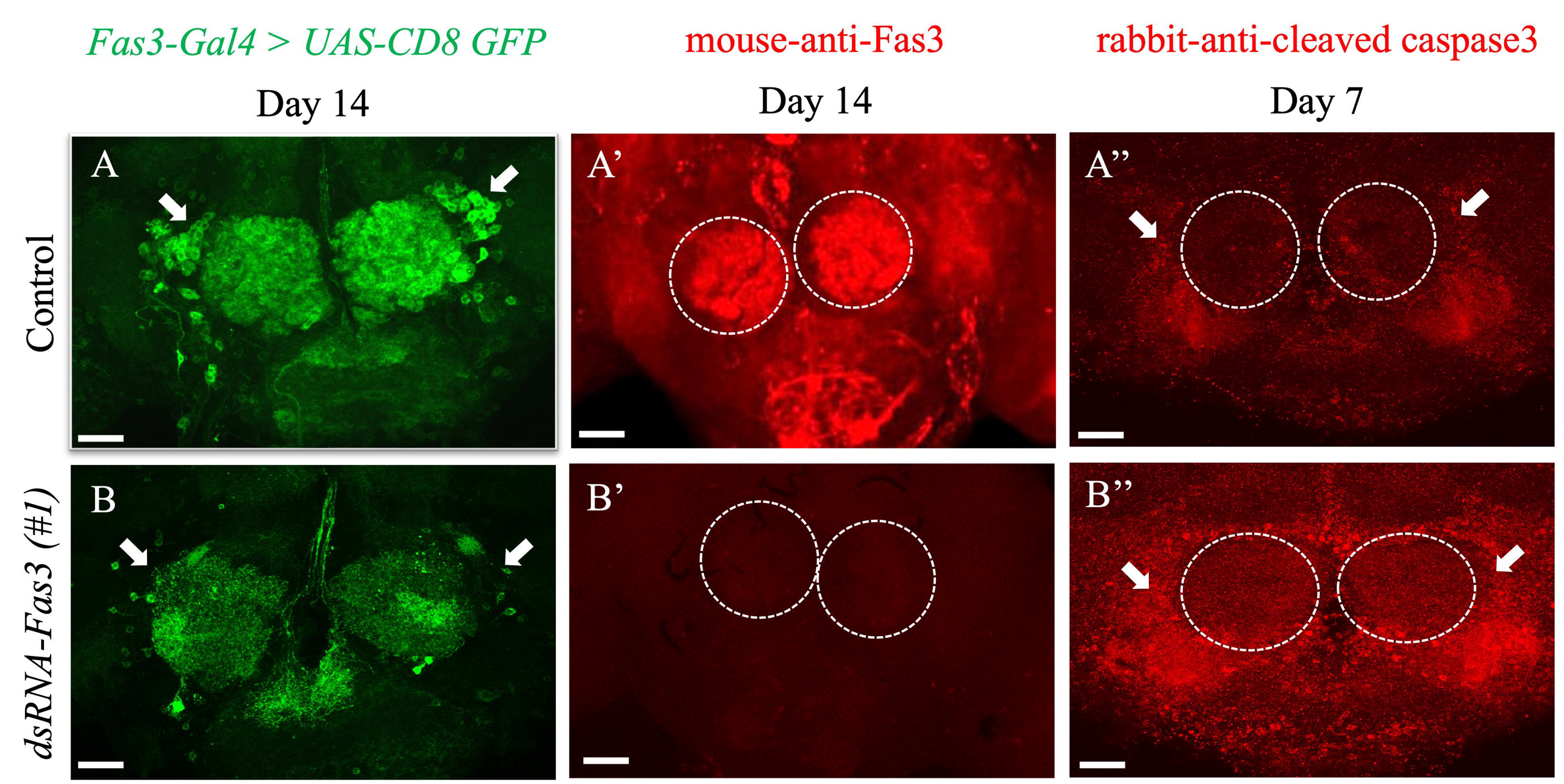
*Fas3* knockdown in all *Fas3*-expressing cells results in widespread death of olfactory neurons. (A, B) Impact of *Fas3* knockdown on olfactory circuit neurons at day 14 post eclosion. The loss of neurons after *Fas3* knockdown is evident from comparing the *UAS-CD8 GFP* labelled neurons in the controls (white arrows in A) and the lack of these neurons in the knockdowns (white arrows in B) (n=10). (A’, B’) Immunostaining using a mouse-anti-Fas3 antibody in controls and knockdowns confirms *Fas3* knockdown by day 14 (n=10). (A”, B”) Cleaved caspase 3 immunolabelling at day 7 confirms that caspase activity is significantly increased in *Fas3* knocked down brains (white arrows in B”) as compared to that in the controls (white arrows in A”) (n=10). The white dotted circles outline the location of the adult olfactory lobes. The scale bars represent 20 μm. Images shown are maximum intensity projections. Genotypes: (A, A’, A”) *UAS-Dicer2; Fas3-Gal4/ UAS-dsRNA-lacZ; tubGal80^ts^/ UAS-CD8 GFP* (B, B’, B”) *UAS-Dicer2; Fas3-Gal4/ UAS-dsRNA-Fas3 (#1); tubGal80^ts^/ UAS-CD8 GFP.* See Supplementary Tables S1-S3 for the additional details on the fly lines.

### Reduced *Fas3* expression results in cell-autonomous cell death

To further confirm that *Fas3* knockdown-mediated cell death occurs in subsets of LNs, we used *LN-Gal4* drivers to restrict Fas3 knockdown to small groups of olfactory interneurons. We conducted this experiment using multiple *UAS-dsRNA-Fas3* lines. Due to the differences in the genetic backgrounds of the *UAS-dsRNA-Fas3* lines, we used a panel of relevant controls from each respective RNAi library. These are detailed in Supplementary Tables S2-S3. The strongest *NP3056-Gal4* mediated knockdown of *Fas3* resulted in a ∼50% loss of *NP3056-Gal4* expressing neurons by day 14 (marked by white arrows in Fig. 5A-G); from 27.15 ± 0.6 neurons in controls to 13.35 ± 0.8 in *Fas3* knockdowns (P ≤ 0.0001) (Fig. 5H, I). A similar cell-autonomous cell death phenotype was also observed when Fas3 was knocked down using other *LN-GAL4* drivers; *GH298-Gal4* (Fig. S2) and *189Y-Gal4* (Fig. S3). Taken together, our data confirm that *Fas3* knockdown results in cell-autonomous death of LNs.

**Fig. 5.**
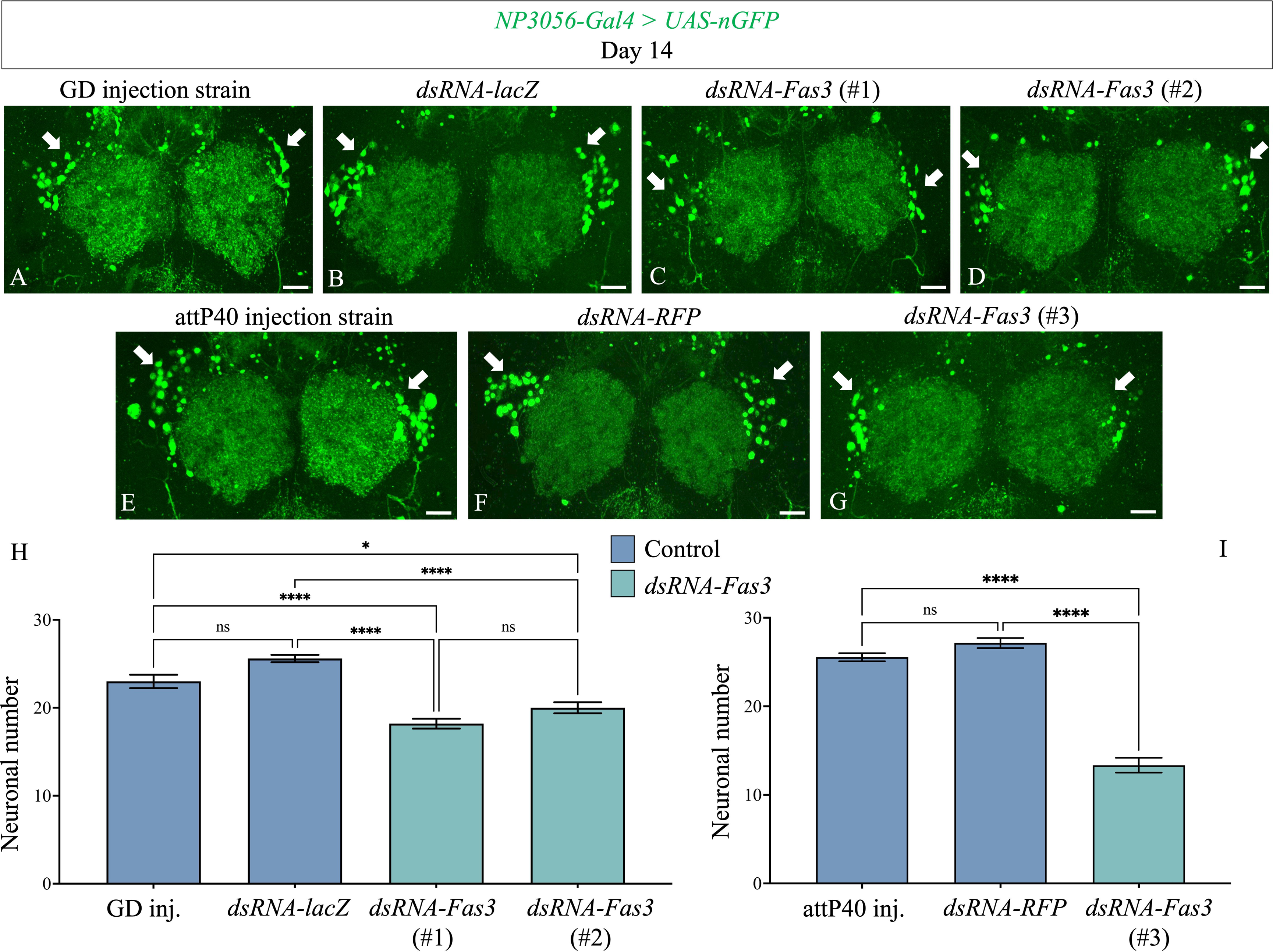
*Fas3* knockdown in a subset of LNs results in restricted neuronal death. (A-D) *NP3056-Gal4* driven knockdown of *Fas3* using the GD RNAi library results in neuronal death by day 14 post eclosion (white arrows). (E-G) Knockdown of *Fas3* using the TRiP RNAi library shows the same impact on LNs (white arrows). The LN neuron subset is labeled with nGFP driven by *NP3056-Gal4* (A-G). The scale bars represent 20 μm. Images shown are maximum intensity projections. (H) Neuronal quantification confirming the reduction of neurons observed in the images A-D. (I) Neuronal quantification confirming the neuronal death observed in the images E-G. Data shown are mean ± SEM. ‘ns’ denotes P > 0.05, ‘*’ denotes P ≤ 0.05, and ‘****’ denotes P ≤ 0.0001 (one-way ANOVA and Tukey’s multiple comparison tests; n=20). Genotypes: (A) *w; tubGal80^ts^, UAS-nGFP/+; NP3056-Gal4/+* (B) *w; tubGal80^ts^, UAS-nGFP/ UAS-dsRNA-lacZ; NP3056-Gal4/+* (C) *UAS-Dicer2; tubGal80^ts^, UAS-nGFP/ UAS-dsRNA-Fas3 (#1); NP3056-Gal4/+* (D) *w; tubGal80^ts^, UAS-nGFP/ UAS-dsRNA-Fas3 (#2); NP3056-Gal4/+* (E) *w; tubGal80^ts^, UAS-nGFP/ attP40; NP3056-Gal4/+* (F) *w; tubGal80^ts^, UAS-nGFP/ UAS-dsRNA-RFP; NP3056-Gal4/+* (G) *w; tubGal80^ts^, UAS-nGFP/ UAS-dsRNA-Fas3 (#3); NP3056-Gal4/+.* See Supplementary Tables S1-S3 for the additional details on the fly lines.

### *Fas3* knockdown-mediated neuronal death is rescued by overexpressing *p35*

To better understand the time course of neuronal loss after *Fas3* knockdown, we used the pan-neuronal driver, *elav-Gal4*, to knockdown *Fas3*. The knockdown was again restricted to the adults using the TARGET system. Here, we examined neuronal numbers at days 3, 5, 7, 10 and 14 after the initiation of Fas3 knockdown (Fig. 6A1-A5, B1-B5). Pan-neuronal knockdown of *Fas3* recapitulated the neuronal death we observed when knocking down *Fas3* in subsets of olfactory neurons. After pan-neuronal Fas3 knockdown, we started to observe the loss of *Fas3* immunoreactivity by day 7, with significant death of olfactory circuit neurons evident by day 10 (Fig. 6A3-A4, B3-B4), which continued through day 14 (Fig. 6A5, B5). These results corroborate our observations using the *Fas3-Gal4* and *LN-Gal4* drivers and add a time course to the observed cell death.

**Fig. 6.**
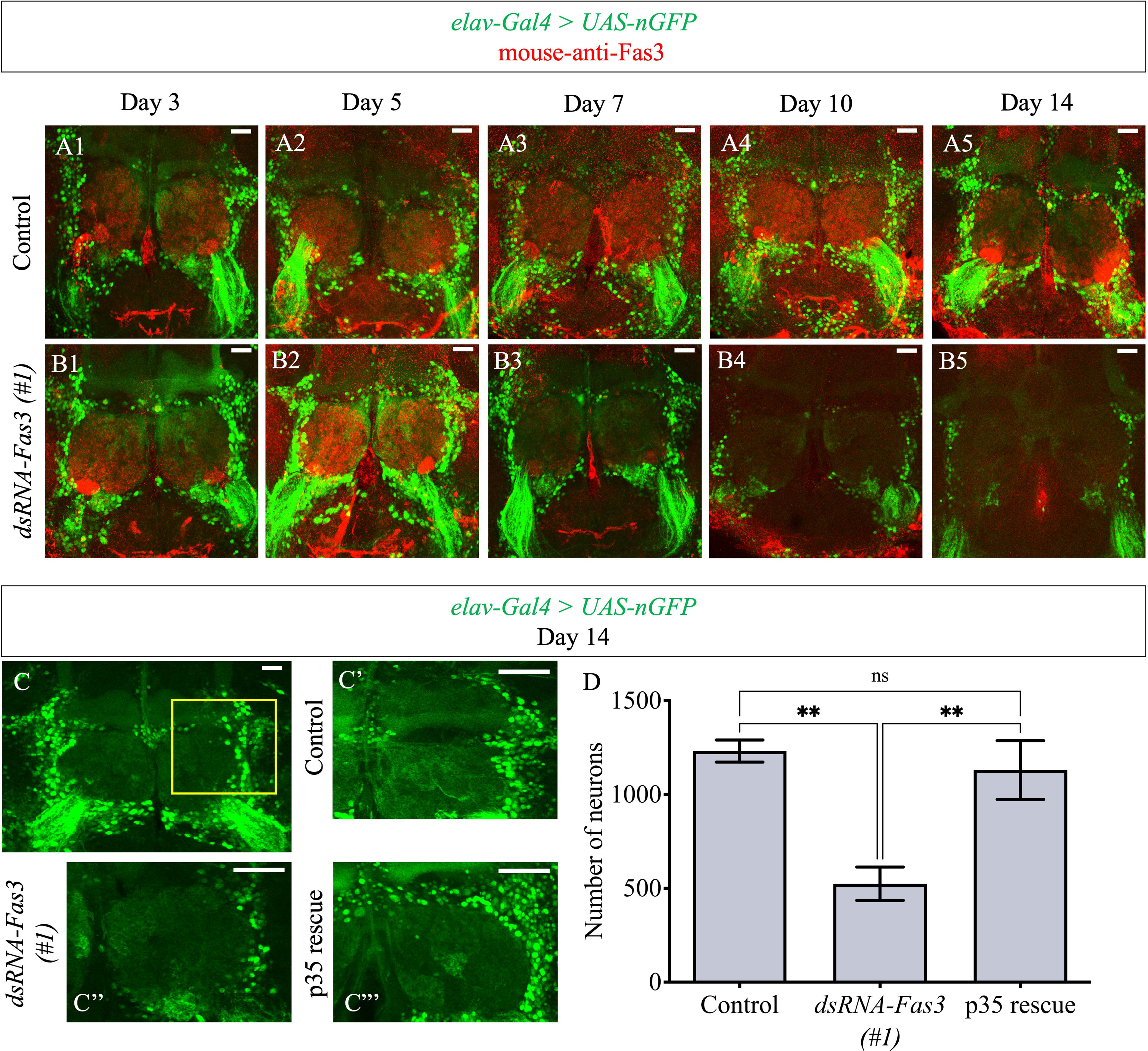
*Fas3* knockdown-mediated neuronal death can be rescued by overexpressing *p35*. (A1-A5, B1-B5) Impact of a pan-neuronal knockdown of *Fas3* on the olfactory circuit as shown from representative images on days 3, 5, 7, 10 and 14 post eclosion. A significant reduction in the neuronal number is evident in the knockdowns on day 10 (A4 vs B4) and day 14 (A5 vs B5). Fas3 immunolabelling is also significantly lower in the knockdowns starting from day 10 (A4 vs B4, A5 vs B5). The knockdown brains from earlier time points do not seem to be any different from the controls (A-A3 vs B1-B3). (C-D) Rescue of *Fas3* knockdown-mediated neuronal death by overexpressing *p35*. (C) A representative image used for quantification. The nuclear GFP driven by *elav-Gal4* in the area surrounding the olfactory lobes (ROI bordered by a yellow box) was quantified. (C’, C”, C”’) Representative images of the ROI used for neuronal quantification. *elav-Gal4* mediated knockdown of *Fas3* results in the loss of LN (C’ vs C”), which is rescued by overexpressing *p35* (C”’). All scale bars represent 20 μm. Images shown are maximum intensity projections. (D) Quantification of neurons in the controls, knockdowns and p35 rescue, confirming a complete rescue of neuronal death. Data shown are mean ± SEM. ‘ns’ denotes P > 0.05, and ‘**’ denotes P ≤ 0.01 (one-way ANOVA and Tukey’s multiple comparison tests; n=20). Genotypes<u>: (</u>A1-A5) *elav-Gal4, UAS-Dicer2;; tubGal80^ts^, UAS-nGFP/+* (B1-B5) *elav-Gal4, UAS-Dicer2; UAS-dsRNA-Fas3 (#1)/+; tubGal80^ts^, UAS-nGFP/+* (C’) *elav-Gal4, UAS-Dicer2;; tubGal80^ts^, UAS-nGFP/+* (C”) *elav-Gal4, UAS-Dicer2; UAS-dsRNA-Fas3 (#1)/+; tubGal80^ts^, UAS-nGFP/+* (C”’) *elav-Gal4, UAS-Dicer2; UAS-dsRNA-Fas3 (#1)/+; tubGal80^ts^, UAS-nGFP/ UAS-p35.* See Supplementary Tables S1-S3 for the additional details on the fly lines.

Since we observed significantly increased cleaved Caspase-3 immunostaining in *Fas3* knockdowns (Fig. 4A”, B”) which is an indicator of apoptosis, we tested if neuronal death can be rescued by blocking the apoptotic pathway. Using *elav-Gal4*, we knocked down Fas3 in the presence of overexpressed p35, an anti-apoptotic protein. We also used *UAS-nGFP* to count neuronal numbers on day 14. We chose the area of LN neurons around each olfactory lobe as the region of interest (ROI) for quantification (indicated by the yellow box in Fig. 6C). Notably, co-expressing *UAS-p35* with *UAS-dsRNA-Fas3* completely rescued the neuronal death phenotype (Fig. 6C-D). The number of neurons (represented by nuclear GFP) in *Fas3* knockdowns was reduced by almost 50% compared to the control and p35 rescue (Fig. 6D). These results confirm that *Fas3* knockdown-mediated neuronal death can be rescued by blocking the apoptotic pathway.

## DISCUSSION

Previous studies have provided insight into the emergence and integration of neurons into the *Drosophila* olfactory circuit; however, very little is understood regarding the maintenance of these neurons in the adult. This study begins to reveal the importance of the stable expression of Fas3, an axon guidance cell adhesion protein, in the maintenance of the adult olfactory circuit. We demonstrated that adult-specific knockdown of *Fas3* leads to the death of local olfactory interneurons which can be rescued by expressing the anti-apoptotic protein p35. Although the dynamic incorporation and function of local interneurons into the rodent olfactory circuit have been examined before (Adam and Mizrahi, 2010; Belluzzi et al., 2003), this is the first study to reveal a critical role for a guidance protein in the maintenance of *Drosophila* interneuron survival. Since we used the TARGET system to restrict the knockdown to the adult, the observed phenotypes are not a result of developmental defects but are due to defects in the maintenance of the adult circuit. To address the potential issue of off-target effects while using RNAi (i.e., knockdown of the expression of multiple genes by the same *dsRNA/ shRNA*), we employed several approaches. First, we picked *UAS-dsRNA* /-*shRNA* lines targeting *Fas3* that were used in previous studies where no off-target effects were reported (Hu et al., 2013). In addition, the observed phenotypes were confirmed using five different Gal4 drivers (*Fas3-Gal4, elav-Gal4, NP3056-Gal4, GH298-Gal4, 189Y-Gal4*) and multiple different *UAS-dsRNA*/-*shRNA* lines and controls (Supplementary Tables S2-S3). This includes *UAS-dsRNA* lines that are integrated into specific attP sites, ensuring that experimental and control *UAS-transgenes* are controlled for position effect (Heigwer et al., 2018).

Though there is evidence for intraglomerular plasticity in the adult *Drosophila* olfactory circuit (Berdnik et al., 2006), this study is the first to show cellular plasticity examine maintenance of the olfactory circuit at the level of a neuronal survival. population., resulting from a cell death pathway that is suppressed by cell adhesion proteins. Cellular plasticity in the adult olfactory system of several species is attributed to their sensory afferents and to subsets of interneurons that are continuously replaced throughout adulthood (Bayramli et al., 2017; Durante et al., 2020; Fernandez-Hernandez et al., 2020 preprint; Graziadei and Okano, 1979; Monti Graziadei and Graziadei, 1979; Weiler and Farbman, 1997). Sustained neurogenesis and apoptosis of sensory neurons has been detected in the antennae of adult *Drosophila*, supporting an ongoing turnover in the olfactory system of adult flies (Fernandez-Hernandez et al., 2020 preprint). In vertebrates, olfactory sensory neuron replacement not only compensates for wear-out processes in the periphery but also contributes to additional glomerular innervations in response to the organism’s experience in adulthood (Jones et al., 2008). In the olfactory epithelium, where mature neurons die and are replaced throughout adult life, tight control over the mechanism of neuronal death is required to avoid tumorigenesis or premature depletion of olfactory sensory neurons. Both *in vivo* and *in vitro* studies have shown that olfactory sensory neurons die through an intrinsically programmed process, involving caspase activity and mediated by jun N-terminal kinase (JNK) signaling (Gangadhar et al., 2008).

In adult rodents, the local circuitry of olfactory interneurons is highly dynamic both at the population level and within individual cells. At the population level, a fraction of rodent interneurons undergo continuous replacement, while at the single-cell level, their dendritic morphology continues to change even after they reach maturity (Adam and Mizrahi, 2010; Belluzzi et al., 2003). The specific subpopulations of interneurons that either remain stable or undergo changes, as well as the extent of these changes, are still not fully understood. However, multiple studies provide compelling evidence for adult neurogenesis of olfactory interneurons in rodents (Kornack and Rakic, 2001; Ming and Song, 2011; Rosselli-Austin and Altman, 1979). Their survival has been shown to be dependent on sensory activity, with odor exposure or olfactory learning promoting neuronal survival (Petreanu and Alvarez-Buylla, 2002; Rochefort et al., 2002). In contrast, there is no evidence for adult neurogenesis of local olfactory interneurons in *Drosophila*. This rules out the possibility that the loss of *Fas3* may be inhibiting the proliferation and differentiation of neural stem cells into local interneurons, thereby resulting in a reduced neuronal number. Moreover, the increased cleaved Caspase 3 labeling in the *Fas3* knockdown brains and the p35 rescue of neuronal death imply that a caspase-dependent cell death pathway is activated in the LNs upon *Fas3* knockdown. Although this may be a newly discovered role for *Fas3*, there is previous evidence associating cell death with the loss of expression of other cell adhesion molecules, such as integrins (Stupack, 2005), and axon guidance genes, such as Semaphorins and Plexins (Vaikakkara Chithran et al., 2023).

*Drosophila* local interneurons (LNs) are diverse in their neurotransmitter profiles, glomerular innervation patterns, and odor response properties. Neuronal subpopulations labelled by *NP3056-, GH298-*, and *189-Gal4* drivers show a patchy glomerular innervation pattern in the olfactory lobe. Each of the *LN-Gal4* drivers label a subset of both larval and adult LNs with at least 189-Gal4 and NP3056-Gal4 showing no overlap of expression (Chou et al., 2010). No two neurons have identical innervation patterns, suggesting that these innervation patterns may be established through cell-cell interactions among LNs (Chou et al., 2010). It may be speculated that the expression of *Fas3*, a homophilic cell adhesion molecule, is required for maintaining the LN-LN interactions in adults which in turn is required for maintaining their patchy innervation. A role for CAMs in maintaining axonal morphologies has been demonstrated previously. For example, it has been shown that JNK signaling is required cell autonomously for axon pruning in mushroom body neurons by negatively regulating the plasma membrane localization of *Fas2*, another cell adhesion molecule of the Ig superfamily (Bornstein et al., 2015). Overexpression of *Fas2* or other CAMs such as *Fas1*, *Fas3* or *Neuroglian* (*Nrg*) was sufficient to inhibit pruning and high levels of CAMs were observed in unpruned axons. Thus, it is possible that *Fas3* knockdown disrupts LN-LN interactions, triggering changes in their axonal/ dendritic morphologies and glomerular innervation, eventually causing neuronal degeneration and death. The importance of maintaining cell-cell interactions is also evident from the aptly called “community effect” observed among precursor cells during development (Saka et al., 2011). Cells in the developing embryo are in constant communication with their neighbors and this interaction is necessary for them to maintain tissue-specific gene expression and differentiate in a coordinated manner. Loss of signaling from neighbors can cause cellular dedifferentiation and death. Similarly, in the adult brain, the loss of cell-cell communication following *Fas3* knockdown may be resulting in neuronal degeneration and death. In addition, *Fas3* is known to genetically interact with other cell adhesion genes including *armadillo* (Greaves et al., 1999), *shotgun* (Greaves et al., 1999), *Toll* (Rose and Chiba, 1999), and *Nrg* (Lu et al., 2014). Ongoing studies will examine if loss of *Fas3* downregulates the expression of other CAMs eventually contributing to cell death.

To maintain survival over the course of their lifetimes, neurons have the ability to adapt certain biological processes to prevent cell death. Cell death pathways are often activated in developing neurons to fine-tune the number of neurons required for the precise formation of neural circuits (Davies, 2003; Kole et al., 2013; Oppenheim, 1991). However, mature neurons promote survival by employing multiple mechanisms to prevent cell death. This shift from permitting cell death to ensuring cellular survival is imperative in mature neurons, as in most cases they must maintain the neuronal circuitry for an organism’s entire lifetime (Benn and Woolf, 2004; Kole et al., 2013). We speculate that maintaining cell-cell interactions by continued expression of guidance cues and cell adhesion proteins molecules such as *Fas3* is a survival mechanism in adult neurons and its loss releases the apoptotic breaks in mature neurons, resulting in cell death. However, the mechanisms downstream of *Fas3* loss that result in death require further detailed analysis. Previous studies have discovered several intracellular signaling cascades that are activated upon the homophilic binding of CAMs (Kiryushko et al., 2004). For example, Leukocyte-antigen-related-like (Lar), a protein that resembles CAMs of the Ig superfamily, controls motor axon guidance in *Drosophila* through protein-tyrosine-phosphatase activity (Krueger et al., 1996). The intracellular region of Fas3 contains two Serine residues that are potential sites of phosphorylation by Protein Kinase C and a Tyrosine residue that may facilitate the attachment of adapter proteins responsible for internalization into coated pits (Snow et al., 1989). Phosphorylation of CAMs such as integrins, N-CAM and L1 has been extensively studied and recognized as a fundamental event in regulating interactions with cytoplasmic proteins to induce alterations in cell adhesion and signaling (Gahmberg and Grönholm, 2022; Matthias and Horstkorte, 2006; Wong et al., 1996). Furthermore, previous research has suggested a role for coated pits in mediating interactions between neuronal growth cones and neighboring cells (Bastiani and Goodman, 1984). Thus, loss of *Fas3* may result in the impairment of cell signaling, endocytosis and protein trafficking, leading to the disruption of cell adhesion and possibly cell death. Further research on intracellular signaling cascades following Fas3 homophilic binding is required to understand how *Fas3* maintains cell survival in LNs.

The main goal of this study was to examine whether *Fas3* has an essential function in the maintenance of adult neural circuits. In addition to the maintenance of adult olfactory neurons, we also demonstrated that *Fas3* is necessary for the survival of adult *Drosophila* which corroborates our previous finding that pan-neuronal expression of axon guidance genes in adults is essential for longevity (Vaikakkara Chithran et al., 2023). The observed reduction in the survival rate upon *Fas3* knockdown could result from the dysfunction of any number of neuronal populations or non neuronal tissues that express Fas3 in the adult *Drosophila* nervous system. Though we have focused on the role of *Fas3* in the adult olfactory circuit in this study, it is important to note that *Fas3* is also expressed in the optic lobes, suboesophageal ganglion, ventral nerve cord and non neuronal tissues. Compromising Fas3 function in multiple neural circuits or none neuronal tissues could result in reduced organismal survival. Examining these additional neuronal populations will determine if *Fas3* is required to maintain their survival during adulthood, similar to its function in adult LNs.

## MATERIALS AND METHODS

### Drosophila genetics

Reducing axon guidance gene expression by RNAi in the adult nervous system was carried out using the TARGET system. Gal4 driver line virgin females (see Supplementary Table S1) were crossed with *UAS-dsRNA*/ *-shRNA* males (see Supplementary Table S2) or the appropriate negative control line males (see Supplementary Table S3). The crosses were set up at 18°C and the progeny (F1) was switched to 29°C immediately after eclosion to repress Gal80^ts^ and activate the Gal4-UAS system. All analyses were performed on day 14 old adult male flies unless otherwise stated.

*Drosophila melanogaster* stocks used in this study were maintained on standard cornmeal food, at 18, 25 or 29°C in environment rooms set at 70% humidity. The stocks used in this study were received from Bloomington *Drosophila* Stock Center (BDSC), Vienna *Drosophila* Resource Center (VDRC) and Kyoto *Drosophila* Stock Center (DGRC) and are listed in Supplementary Tables S1-S4.

### Survival assay

F1 progeny males were collected immediately after eclosion and switched to 29°C. They were maintained in vials of 10 and their survival was recorded every day for the next 20 days. Flies were flipped into fresh vials every week. Multiple *UAS-dsRNA/-shRNA* lines and controls were tested and for each line, 3 vials containing 10 flies were assayed. All results were analyzed using two-way ANOVA and Tukey’s multiple comparison tests using GraphPad Prism.

### Immunohistochemistry and imaging

Adult male *Drosophila* brains and VNCs were dissected in ice-cold 1x PBS, over a period of 30 minutes. The tissue was fixed, on poly-Lysine coated slides, in 4% PFA at room temperature (RT) for 30 minutes followed by 3x 5 minutes washes with 0.1% PBT. The samples were blocked in 5% PBTN for 2 hours at RT, and then incubated in primary antibodies overnight at 4°C. The next day, samples were washed 3x 30 minutes in 0.1% PBT. They were then blocked in 5% PBTN for 1 hour at RT and incubated in secondary antibody for 3 hours at RT in the dark. For each experiment, controls and knockdowns were dissected and labelled in parallel. The samples were washed 3x 30 minutes in 0.1% PBT. They were mounted in Vectashield mounting media (Vector Laboratories Inc.), coverslipped and sealed with nail polish. Primary antibodies used in this study were as follows: mouse-anti-Bruchpilot nc82 (DSHB, 1:20), mouse-anti-Fasciclin III 7G10 (DSHB, 1:400), mouse-anti-β-gal (DSHB, 1:50), chicken-anti-β-gal (Abcam, 1:1000), rabbit-anti-cleaved caspase 3 (Asp175) (Cell Signaling Technology, 1:300). Secondary antibodies used in this study were as follows: donkeyanti-chicken cy3 (Jackson, 1:250) donkey-anti-mouse cy3 (Jackson, 1:250), donkey-anti-rabbit cy3 (Jackson, 1:250). All images were acquired using the Zeiss LSM 880 Fast Airyscan inverted confocal microscope in a Z-stack. Images were processed using the ‘Airyscan processing’ feature using the Zen-black software. The images were acquired in multiple tiles and stitched post Airyscan processing. The images included in this study are maximum-intensity projections of representative images and all images were acquired with the same parameters.

### Neuronal quantification and Statistical analysis

In *elav-Gal4* driven experiments, the area around the olfactory lobe was selected as the region of interest and neurons (represented by nlsGFP) were quantified using a MATLAB program kindly provided by Katie Goodwin (University of British Columbia). In *LN-Gal4* driven experiments, the olfactory circuits neurons were quantified manually in a 3D-model (using Zen-blue image processing software). The results were analysed using one-way ANOVA and Tukey’s multiple comparison tests using GraphPad Prism.

## Acknowledgements

The authors extend sincere thanks to Katie Goodwin for kindly sharing the MATLAB program for image analysis. The antibodies used in this study were obtained from Developmental Studies Hybridoma Bank (DSHB), Abcam, Cell Signaling Technology and Jackson ImmunoResearch Laboratories Inc. *Drosophila melanogaster* stocks were received from Bloomington *Drosophila* Stock Center (BDSC), Vienna *Drosophila* Resource Center (VDRC) and Kyoto *Drosophila* Stock Center (DGRC).

## Author contributions

Conceptualization: A.V.C, D.W.A, T.OC; Methodology: A.V.C; Validation: A.V.C, D.W.A, T.OC; Formal analysis: A.V.C; Investigation: A.V.C; Writing-original draft: A.V.C; Writing-review & editing: A.V.C, D.W.A, T.OC; Visualization: A.V.C; Supervision: D.W.A, T.OC; Funding acquisition: A.V.C, T.OC

## Data availability

The datasets generated during and/ or analyzed during the current study are available from the corresponding author on request.

## References

Adam, Y. and Mizrahi, A. (2010). Circuit formation and maintenance—perspectives from the mammalian olfactory bulb. Current opinion in neurobiology 20, 134–140.

Bastiani, M. J. and Goodman, C. S. (1984). Neuronal growth cones: specific interactions mediated by filopodial insertion and induction of coated vesicles. Proceedings of the National Academy of Sciences 81, 1849–1853.

Bayramli, X., Kocagöz, Y., Sakizli, U. and Fuss, S. H. (2017). Patterned arrangements of olfactory receptor gene expression in zebrafish are established by radial movement of specified olfactory sensory neurons. Scientific Reports 7, 5572.

Belluzzi, O., Benedusi, M., Ackman, J. and LoTurco, J. J. (2003). Electrophysiological differentiation of new neurons in the olfactory bulb. Journal of Neuroscience 23, 10411–10418.

Benn, S. C. and Woolf, C. J. (2004). Adult neuron survival strategies-slamming on the brakes. Nature Reviews Neuroscience 5, 686–700.

Berdnik, D., Chihara, T., Couto, A. and Luo, L. (2006). Wiring stability of the adult Drosophila olfactory circuit after lesion. Journal of Neuroscience 26, 3367–3376.

Bornstein, B., Zahavi, E.E., Gelley, S., Zoosman, M., Yaniv, S.P., Fuchs, O., Porat, Z., Perlson, E. and Schuldiner, O. (2015). Developmental axon pruning requires destabilization of cell adhesion by JNK signaling. Neuron 88, 926–940.

Cajal, S. (1890). Sobre la aparici.n de las expansiones celulares en la m.dula embrionaria. Gac.Sanit. Barc. 2, 413–419.

Chiba, A., Snow, P., Keshishian, H. and Hotta, Y. (1995). Fasciclin III as a synaptic target recognition molecule in Drosophila. Nature 374, 166–168.

Chou, Y. H., Spletter, M. L., Yaksi, E., Leong, J. C., Wilson, R. I. and Luo, L. (2010). Diversity and wiring variability of olfactory local interneurons in the Drosophila antennal lobe. Nature neuroscience 13, 439–449.

Couto, A., Alenius, M. and Dickson, B. J. (2005). Molecular, anatomical, and functional organization of the Drosophila olfactory system. Current Biology 15, 1535–1547.

Das, S., Sadanandappa, M.K., Dervan, A., Larkin, A., Lee, J.A., Sudhakaran, I.P., Priya, R., Heidari, R., Holohan, E.E., Pimentel, A. et al. (2011). Plasticity of local GABAergic interneurons drives olfactory habituation. Proceedings of the National Academy of Sciences 108, E646–E654.

Davies, A. M. (2003). Regulation of neuronal survival and death by extracellular signals during development. The EMBO journal 22, 2537–2545.

Durante, M.A., Kurtenbach, S., Sargi, Z.B., Harbour, J.W., Choi, R., Kurtenbach, S., Goss, G.M., Matsunami, H. and Goldstein, B.J. (2020). Single-cell analysis of olfactory neurogenesis and differentiation in adult humans. Nature neuroscience 23, 323–326.

Eddison, M., Belay, A. T., Sokolowski, M. B. and Heberlein, U. (2012). A genetic screen for olfactory habituation mutations in Drosophila: analysis of novel foraging alleles and an underlying neural circuit. PLoS One 7, e51684.

Elkins, T., Zinn, K., McAllister, L., HoffMann, F. M. and Goodman, C. S. (1990). Genetic analysis of a Drosophila neural cell adhesion molecule: interaction of fasciclin I and Abelson tyrosine kinase mutations. Cell 60, 565–575.

Fernández-Hernández, I., Hu, E. and Bonaguidi, M. A. (2020). Olfactory neuron turnover in adult Drosophila. bioRxiv, 2020-11.

Fishilevich, E. and Vosshall, L. B. (2005). Genetic and functional subdivision of the Drosophila antennal lobe. Current Biology 15, 1548–1553.

Gahmberg, C. G. and Grönholm, M. (2022). How integrin phosphorylations regulate cell adhesion and signaling. Trends in Biochemical Sciences 47, 265–278.

Gangadhar, N. M., Firestein, S. J. and Stockwell, B. R. (2008). A novel role for jun N-terminal kinase signaling in olfactory sensory neuronal death. Molecular and Cellular Neuroscience 38, 518–525.

Graveley, B.R., Brooks, A.N., Carlson, J.W., Duff, M.O., Landolin, J.M., Yang, L., Artieri, C.G., Van Baren, M.J., Boley, N., Booth, B.W. et al. (2011). The developmental transcriptome of Drosophila melanogaster. Nature 471, 473–479.

Graziadei, P. P. C. and Okano, M. (1979). Neuronal degeneration and regeneration in the olfactory epithelium of pigeon following transection of the first cranial nerve. Cells Tissues Organs 104, 220–236.

Greaves, S., Sanson, B., White, P. and Vincent, J. P. (1999). A screen for identifying genes interacting with armadillo, the Drosophila homolog of β-catenin. Genetics 153, 1753–1766.

Harrelson, A. L. and Goodman, C. S. (1988). Growth cone guidance in insects: fasciclin II is a member of the immunoglobulin superfamily. Science 242, 700–708.

Heigwer, F., Port, F. and Boutros, M. (2018). RNA interference (RNAi) screening in Drosophila. Genetics 208, 853–874.

Hu, Y., Roesel, C., Flockhart, I., Perkins, L., Perrimon, N. and Mohr, S. E. (2013). UP-TORR: online tool for accurate and Up-to-Date annotation of RNAi Reagents. Genetics 195, 37–45.

Imai, T., Sakano, H. and Vosshall, L. B. (2010). Topographic mapping-the olfactory system. Cold Spring Harbor perspectives in biology 2, a001776.

Jones, S. V., Choi, D. C., Davis, M. and Ressler, K. J. (2008). Learning-dependent structural plasticity in the adult olfactory pathway. Journal of Neuroscience 28, 13106–13111.

Kiryushko, D., Berezin, V. and Bock, E. (2004). Regulators of neurite outgrowth: role of cell adhesion molecules. Annals of the New York Academy of Sciences 1014, 140–154.

Kole, A. J., Annis, R. P. and Deshmukh, M. (2013). Mature neurons: equipped for survival. Cell death & disease 4, e689–e689.

Kolodkin, A. L. and Tessier-Lavigne, M. (2011). Mechanisms and molecules of neuronal wiring: a primer. Cold Spring Harbor perspectives in biology 3, a001727.

Kornack, D. R. and Rakic, P. (2001). The generation, migration, and differentiation of olfactory neurons in the adult primate brain. Proceedings of the National Academy of Sciences 98, 4752–4757.

Kose, H., Rose, D., Zhu, X. and Chiba, A. (1997). Homophilic synaptic target recognition mediated by immunoglobulin-like cell adhesion molecule Fasciclin III. Development 124, 4143–4152.

Krueger, N. X., Van Vactor, D., Wan, H. I., Gelbart, W. M., Goodman, C. S. and Saito, H. (1996). The transmembrane tyrosine phosphatase DLAR controls motor axon guidance in Drosophila. Cell 84, 611–622.

Laissue, P. P., Reiter, C. H., Hiesinger, P. R., Halter, S., Fischbach, K. F. and Stocker, R. F. (1999). ThreeLdimensional reconstruction of the antennal lobe in Drosophila melanogaster. Journal of Comparative Neurology 405, 543–552.

Larkin, A., Karak, S., Priya, R., Das, A., Ayyub, C., Ito, K., Rodrigues, V. and Ramaswami, M. (2010). Central synaptic mechanisms underlie short-term olfactory habituation in Drosophila larvae. Learning & memory 17, 645–653.

Lin, D. M. and Goodman, C. S. (1994). Ectopic and increased expression of Fasciclin II alters motoneuron growth cone guidance. Neuron 13, 507–523.

Lin, D. M., Fetter, R. D., Kopczynski, C., Grenningloh, G. and Goodman, C. S. (1994). Genetic analysis of Fasciclin II in Drosophila: defasciculation, refasciculation, and altered fasciculation. Neuron 13, 1055–1069.

Liou, N.F., Lin, S.H., Chen, Y.J., Tsai, K.T., Yang, C.J., Lin, T.Y., Wu, T.H., Lin, H.J., Chen, Y.T., Gohl, D.M., et al. (2018). Diverse populations of local interneurons integrate into the Drosophila adult olfactory circuit. Nature communications 9, 2232.

Liu, W. W. and Wilson, R. I. (2013). Glutamate is an inhibitory neurotransmitter in the Drosophila olfactory system. Proceedings of the National Academy of Sciences 110, 10294–10299.

Lledo, P. M., Gheusi, G. and Vincent, J. D. (2005). Information processing in the mammalian olfactory system. Physiological reviews 85, 281–317.

Lu, C. S., Zhai, B., Mauss, A., Landgraf, M., Gygi, S. and Van Vactor, D. (2014). MicroRNA-8 promotes robust motor axon targeting by coordinate regulation of cell adhesion molecules during synapse development. Philosophical Transactions of the Royal Society B: Biological Sciences 369, 20130517.

Matthias, S. and Horstkorte, R. (2006). Phosphorylation of the neural cell adhesion molecule on serine or threonine residues is induced by adhesion or nerve growth factor. Journal of neuroscience research 84, 142–150.

Ming, G. L. and Song, H. (2011). Adult neurogenesis in the mammalian brain: significant answers and significant questions. Neuron 70, 687–702.

Monti Graziadei, G. A. and Graziadei, P. P. C. (1979). Neurogenesis and neuron regeneration in the olfactory system of mammals. II. Degeneration and reconstitution of the olfactory sensory neurons after axotomy. Journal of neurocytology 8, 197–213.

Moreland, T. and Poulain, F. E. (2022). To stick or not to stick: The multiple roles of cell adhesion molecules in neural circuit assembly. Frontiers in Neuroscience 16, 552.

Oppenheim, R. W. (1991). Cell death during development of the nervous system. Annual review of neuroscience 14, 453–501.

Patel, N. H., Snow, P. M. and Goodman, C. S. (1987). Characterization and cloning of fasciclin III: a glycoprotein expressed on a subset of neurons and axon pathways in Drosophila. Cell 48, 975–988.

Petreanu, L. and Alvarez-Buylla, A. (2002). Maturation and death of adult-born olfactory bulb granule neurons: role of olfaction. Journal of Neuroscience 22, 6106–6113.

Pollerberg, G. E., Thelen, K., Theiss, M. O. and Hochlehnert, B. C. (2013). The role of cell adhesion molecules for navigating axons: density matters. Mechanisms of development 130, 359–372.

Rochefort, C., Gheusi, G., Vincent, J. D. and Lledo, P. M. (2002). Enriched odor exposure increases the number of newborn neurons in the adult olfactory bulb and improves odor memory. Journal of Neuroscience 22, 2679–2689.

Rose, D. and Chiba, A. (1999). A single growth cone is capable of integrating simultaneously presented and functionally distinct molecular cues during target recognition. Journal of Neuroscience 19, 4899–4906.

Rosselli-Austin, L. and Altman, J. (1979). The postnatal development of the main olfactory bulb of the rat. Journal of developmental physiology 1, 295–313.

Rutishauser, U. (1993). Adhesion molecules of the nervous system. Current opinion in neurobiology 3, 709–715.

Sadanandappa, M.K., Redondo, B.B., Michels, B., Rodrigues, V., Gerber, B., VijayRaghavan, K., Buchner, E. and Ramaswami, M. (2013). Synapsin function in GABA-ergic interneurons is required for short-term olfactory habituation. Journal of Neuroscience 33, 16576–16585.

Saka, Y., Lhoussaine, C., Kuttler, C., Ullner, E. and Thiel, M. (2011). Theoretical basis of the community effect in development. BMC systems biology 5, 1–14.

Seki, Y., Rybak, J., Wicher, D., Sachse, S. and Hansson, B. S. (2010). Physiological and morphological characterization of local interneurons in the Drosophila antennal lobe. Journal of neurophysiology 104, 1007–1019.

Snow, P. M., Bieber, A. J. and Goodman, C. S. (1989). Fasciclin III: a novel homophilic adhesion molecule in Drosophila. Cell 59, 313–323.

Stocker, R. F., Lienhard, M. C., Borst, A. and Fischbach, K. F. (1990). Neuronal architecture of the antennal lobe in Drosophila melanogaster. Cell and tissue research 262, 9–34.

Stupack, D. G. (2005). Integrins as a distinct subtype of dependence receptors. Cell Death & Differentiation 12, 1021–1030.

Sudhakaran, I. P., Holohan, E. E., Osman, S., Rodrigues, V., Vijayraghavan, K. and Ramaswami, M. (2012). Plasticity of recurrent inhibition in the Drosophila antennal lobe. Journal of Neuroscience 32, 7225–7231.

Tanaka, N. K., Endo, K. and Ito, K. (2012). Organization of antennal lobeLassociated neurons in adult Drosophila melanogaster brain. Journal of Comparative Neurology 520, 4067–4130.

Tanaka, N. K., Ito, K. and Stopfer, M. (2009). Odor-evoked neural oscillations in Drosophila are mediated by widely branching interneurons. Journal of Neuroscience 29, 8595–8603.

Tessier-Lavigne, M. and Goodman, C.S. (1996). The molecular biology of axon guidance. Science 274, 1123–1133.

Twick, I., Lee, J. A. and Ramaswami, M. (2014). Olfactory habituation in Drosophila-odor encoding and its plasticity in the antennal lobe. Progress in Brain Research 208, 3–38.

Vaikakkara Chithran, A., Allan, D. W. and O’Connor, T. P. (2023). Adult expression of Semaphorins and Plexins is essential for motor neuron survival. Scientific Reports 13, 5894.

Weiler, E. and Farbman, A. I. (1997). Proliferation in the rat olfactory epithelium: age-dependent changes. Journal of Neuroscience 17, 3610–3622.

Wong, E. V., Schaefer, A. W., Landreth, G. and Lemmon, V. (1996). Casein kinase II phosphorylates the neural cell adhesion molecule L1. Journal of neurochemistry 66, 779–786.

Yaron, A. and Zheng, B. (2007). Navigating their way to the clinic: emerging roles for axon guidance molecules in neurological disorders and injury. Developmental neurobiology 67, 1216–1231.

